# Structure and inhibitor binding characterization of oncogenic MLLT1 mutants

**DOI:** 10.1101/2021.02.08.430291

**Authors:** Xiaomin Ni, Allyn T. Londregan, Dafydd R. Owen, Stefan Knapp, Apirat Chaikuad

## Abstract

Dysfunction of YEATS-domain-containing MLLT1, an acetyl/acyl-lysine dependent epigenetic reader domain, has been implicated in the development of aggressive cancers. Mutations in the YEATS domain have been recently reported as a cause of MLLT1 aberrant reader function. However, structural basis for the reported alterations in affinity for acetyled/acylated histone has remained elusive. Here, we report the crystal structures of both insertion and substitution present in cancer, revealing significant conformational changes of the YEATS-domain loop 8. Structural comparison demonstrates that such alteration not only altered the binding interface for acetylated/acylated histones, but the sequence alterations in the T1 loop may enable dimeric assembly consistent inducing self-association behavior. Nevertheless, we show that also the MLLT1 mutants can be targeted by developed acetyllysine mimetic inhibitors with affinities similarly to wild type. Our report provides a structural basis for the altered behaviors and potential strategy for targeting oncogenic MLLT1 mutants.

---

Mixed-lineage leukemia translocated to 1 (MLLT1), known also as eleven–nineteen leukemia (ENL), is a key player in epigenetics signaling acting as an Nε-acetylated lysine reader ^*1-4*^. This function of MLLT1 occurs via its YEATS (Yaf9, ENL, AF9, Taf14, and Sas5) domain^*5*^, which in human also exists in other three proteins, including YEATS2, MLLT3 (known also as AF9), and glioma amplified sequence 41 (GAS41 or YEATS4). Besides an ability to recognize acetylation mark, the human YEATS protein module has been shown to exhibit also broader reader activity for other lysine modifications, including propionylation, butyrylation, crotonylation and succinylation^*3, 6-11*^. The reader activities of MLLT1 as well as the other three human YEATS-domain-containing proteins are well established for epigenetics marks encoded on the histone tails, of which the interaction leads to their localization to chromatin and consequently initiates expression of target genes^*8, 12*^. This highlights therefore the important regulatory function of MLLT1 and the YEATS module in transcription.

Molecular mechanisms of recognition of acetylated histone by YEATS domain have been well studied^*2-4, 9, 10, 13*^. Acetyllysine (Kac) and other acylated lysine, such as crotonyllysine (Kcr), bind within the pocket that situates on top of an elongated β-sheet core and is formed largely by the highly conserved aromatic triad (Phe28, Phe59 and Phe78 in MLLT1) from loop 1, 4 and 6, which contributes π-π interactions ^*1-3, 9, 10, 14*^. The binding of Kac acts primarily as an anchor point, while other parts of histone tail are tethered to the surface of YEAST domain along the β-sheet core and loop 8 that is located adjacent to the acetyllysine binding pocket. Despite limited direct contacts, the histone residues that flank N- and C-termini of the central Kac likely determine specificity, which differs among four YEATS-domain-containing proteins^*6*^.

Dysfunction of YEATS-domain-containing proteins has been demonstrated as a factor driving the development of several diseases, essentially cancer^*6*^. MLLT1 or MLLT3 are frequent fusion partners with the human mixed lineage leukemia (MLL), resulting in an oncoprotein can cause aggressive cancer growth^*1, 15, 16*^. Thus, targeting their YEATS protein modules with small molecules has been proposed as a potential chemotherapeutic strategy for treatment of these diseases. Early fragment screening campaigns demonstrated that the Kac binding pocket of MLLT1/3 YEATS domain is druggable with a number of small molecule binders that have been identified ^*17, 18*^. To date, several of potent and selective MLLT1/3 inhibitors have been developed that can serve as chemical probes studying MLLT1/3 fundtion^*19-21*^.

Recently, mutations within YEATS domain have been discovered as another cause of aberrant function of MLLT1. To date, eight MLLT1 mutants have been identified, essentially in Wilms tumors. The mutations are primarily located in loop 8 and can be classified either as an in-frame insertion or substitution with deletion (Figure 1A)^*22, 23*^. Among them, p.117_118insNHL (known also and hereafter referred to as T1), p.112_114PPV> L (known also and hereafter referred to as T2) and p.111_113NPP> K (known also and hereafter referred to as T3) mutants have been most studied due to their high frequencies in cancers^*22, 23*^. Different characteristics of these three mutants compared to wild type have been reported, including weaker affinities for acetylated histone tails^*22, 23*^. In addition, another intriguing property, noted especially for T1, is the ability of the mutant to self-associate, increase occupancy on chromatin resulting in increased recruitment of the super elongation complex (SEC) and as a consequence, boosting the transcription of several oncogenes, such as MYC and HOX^*22*^.

**Figure 1.**
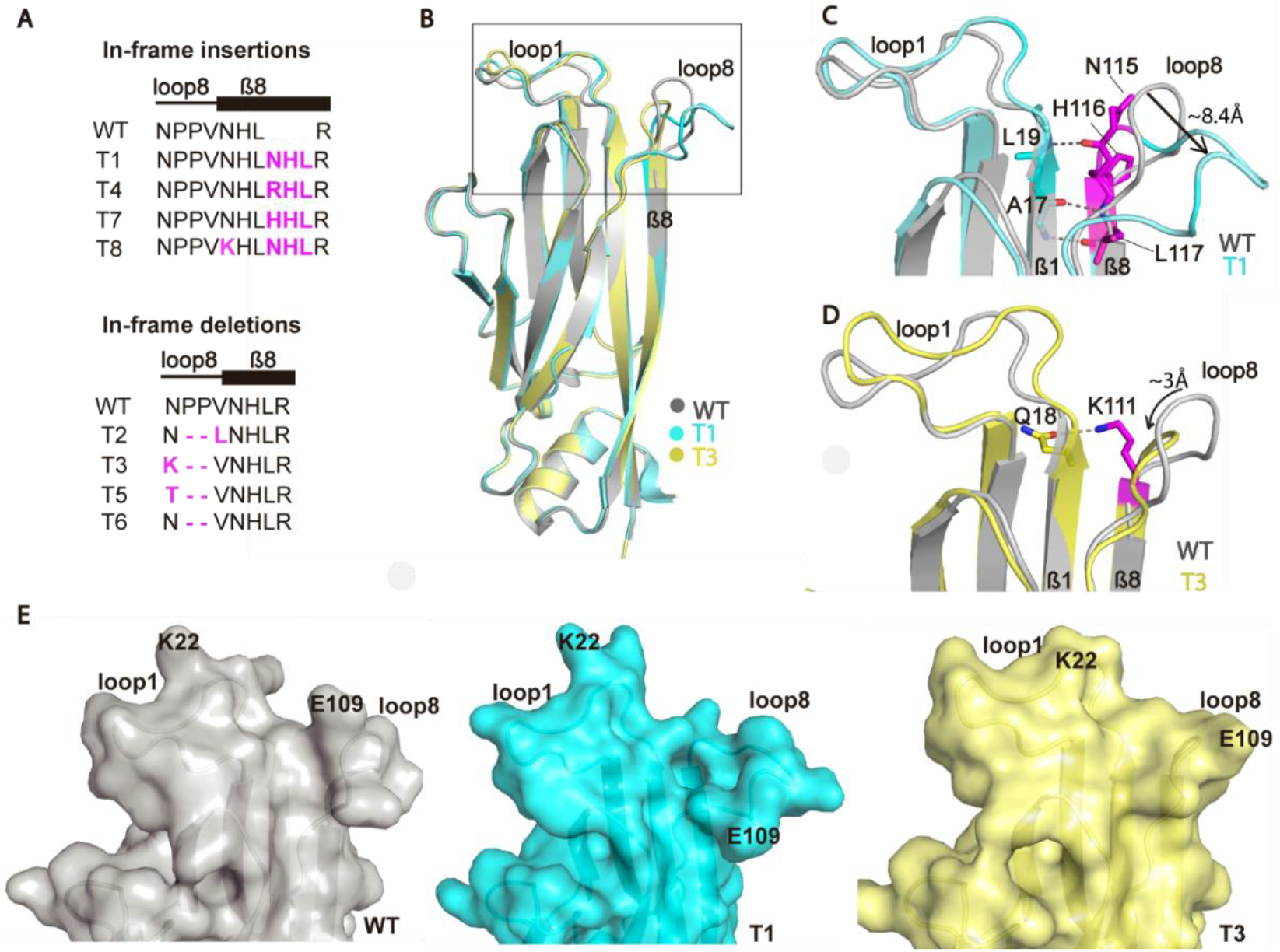
Structural comparison between wild type MLLT1 and T1 and T3 mutants. (A) Eight oncogenic mutations on loop 8 region of MLLT1 (T1 to T8). (B) Superimposition of T1 (cyan), T3 (yellow) and wild type (WT, gray; PDB ID: 6HQ0) structures demonstrates significant alterations of loop 8 caused by insertion and substitution with deletion mutations in T1 and T3, respectively. Close-up reveals an opening and a close conformation of loop 8 in T1 (C) and T3 (D), respectively. (E) Surface representation demonstrates the consequences of the mutations on significant alterations of the protein surface.

In order to provide high resolution models describing the structural consequence of the oncogenic mutations in MLLT1 YEATS domain, we determined the crystal structures of T1 and T3 mutants in complex with an inhibitor (Table 1), providing insights of structural consequences caused by insertions and deletions in YEATS domains. The structures of T1 and T3 were determined to high resolution of ∼1.9-2.1 Å and all elements, essentially the mutated regions, were well-defined by electron density. Structural comparison with wild type^*18*^ revealed that while no significant change was observed for the β-sheet central core and the acetylated- or acylated-lysine binding groove, substantial structural alterations were evident for loop 8 that forms a part of the histone interaction interface (Figure 1B). In T1, the additional three Asn-His-Leu residues (NHL) formed a β-sheet structure, resulting in an extension of both β8 strand and loop 8 that for the latter expanded outward from the core β-sheet with an ∼8.4-Å positional shift in comparison to wild type (Figure 1C). This conformational change created a vast opening channel between loop 1 and 8, which was restricted to only a small groove in wild type. On a contrary, the substitution of Asn-Pro-Pro (aa 111-113) to a single lysine residue shortened and rigidified loop 8 in T3, which was instead packed closer to the β-sheet core (Figure 1D). Such alteration diminished the groove constructed between loop 1 and 8 that otherwise existed in wild type and T1 (Figure 1E). Overall, these crystal structures demonstrated that both types of mutations induced significant conformational changes of loop 8, causing dramatic alterations on the protein surface, albeit with different consequences.

**Table 1.**
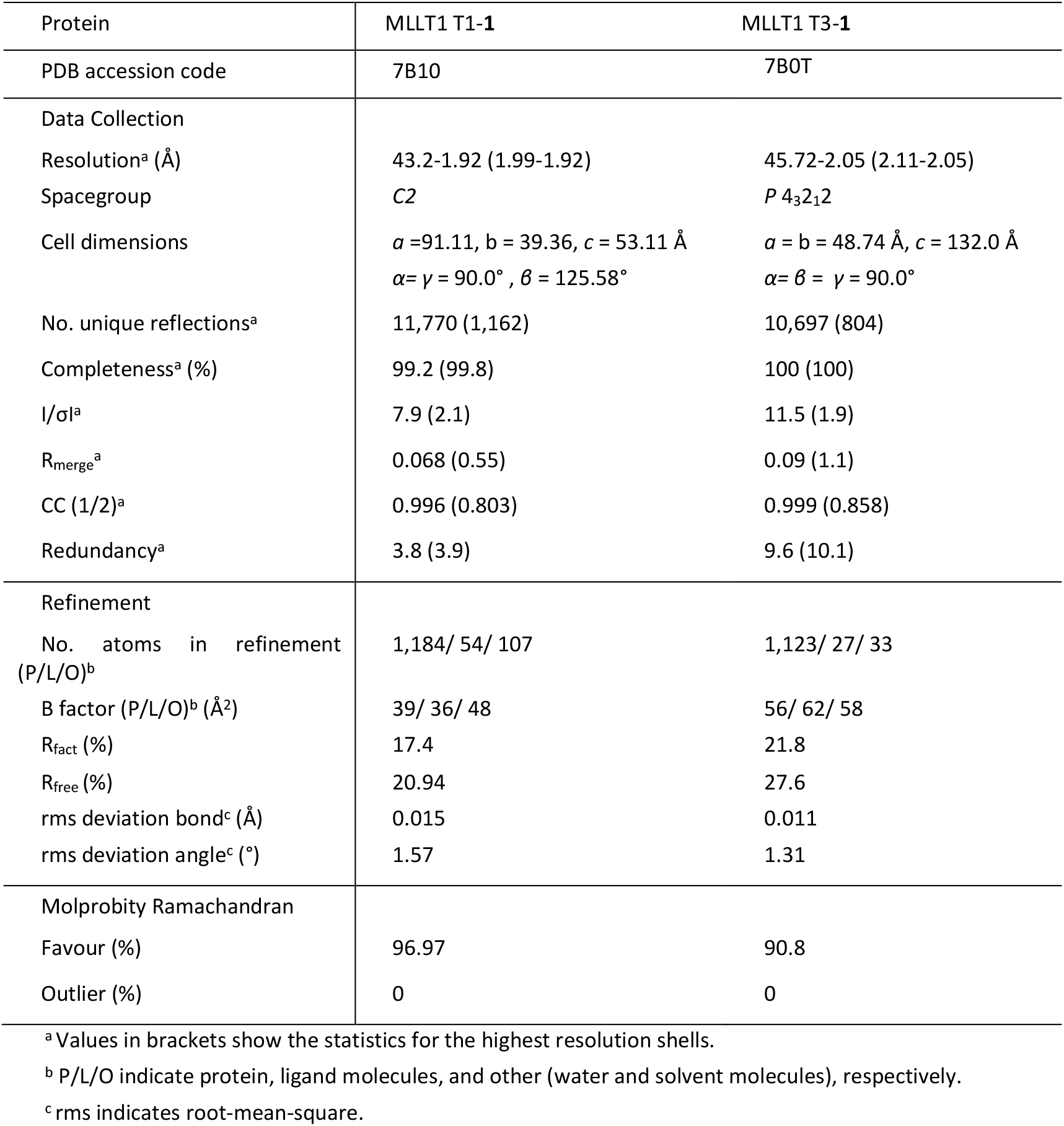
Data collection and refinement statistics

Although the integrity of the acetyl/acyllysine binding pocket remained unaffected, several lines of evidence have shown that the mutations weakened the binding of the acetylated and acylated histone peptides^*22, 23*^. To provide an explanation for such behaviors, we performed structural comparison with the published MLLT1 or MLLT3 (AF9) complexed with well-defined acetylated or acylated histone peptides^*1, 3, 9*^. Comparative structural analyses revealed that while the binding of Kac and Kcr is likely preserved, the alterations of loop 8 could significantly perturb the interactions with the histone N-terminal region (Figure2). In T1, the extended loop 8 the histone docking site, explaining the previously observed ∼3-4-fold weaker affinity for H3K27ac/cr or no detectable binding for H3K9ac in T1 (Figure 2B)^*22, 23*^. In contrast, based on our structural models the strongly negative influence on the binding of histone marks would be less pronounced in T3, in which the binding interface remained mostly unperturbed. Nevertheless, some steric clashes, such as between loop 8 Leu108 and the histone residue at the -3 position from Kac/Kcr, might be anticipated (Figure 2C), providing a potential explanation for subtly lower affinity for H3K27ac/cr or differences in thermodynamic binding behavior for H3K9ac, of which the affinity remained unaffected, in T3 ^*22, 23*^. In addition, our structural comparison and isothermal calorimetry (ITC) data further suggested that significant alterations of the peptide interfaces caused by the mutations will modulate also the interactions with other histone marks, such as at Lys14 and Lys18 (Figure 2). Overall, our comparative structural analyses suggested that although the acetyl/acyllysine binding pocket remains largely unperturbed, different binding conformations of acetylated/acylated histone tail interacting to these oncogenic mutants would likely be expected due an alteration of the peptide binding interface caused by the mutation.

**Figure 2.**
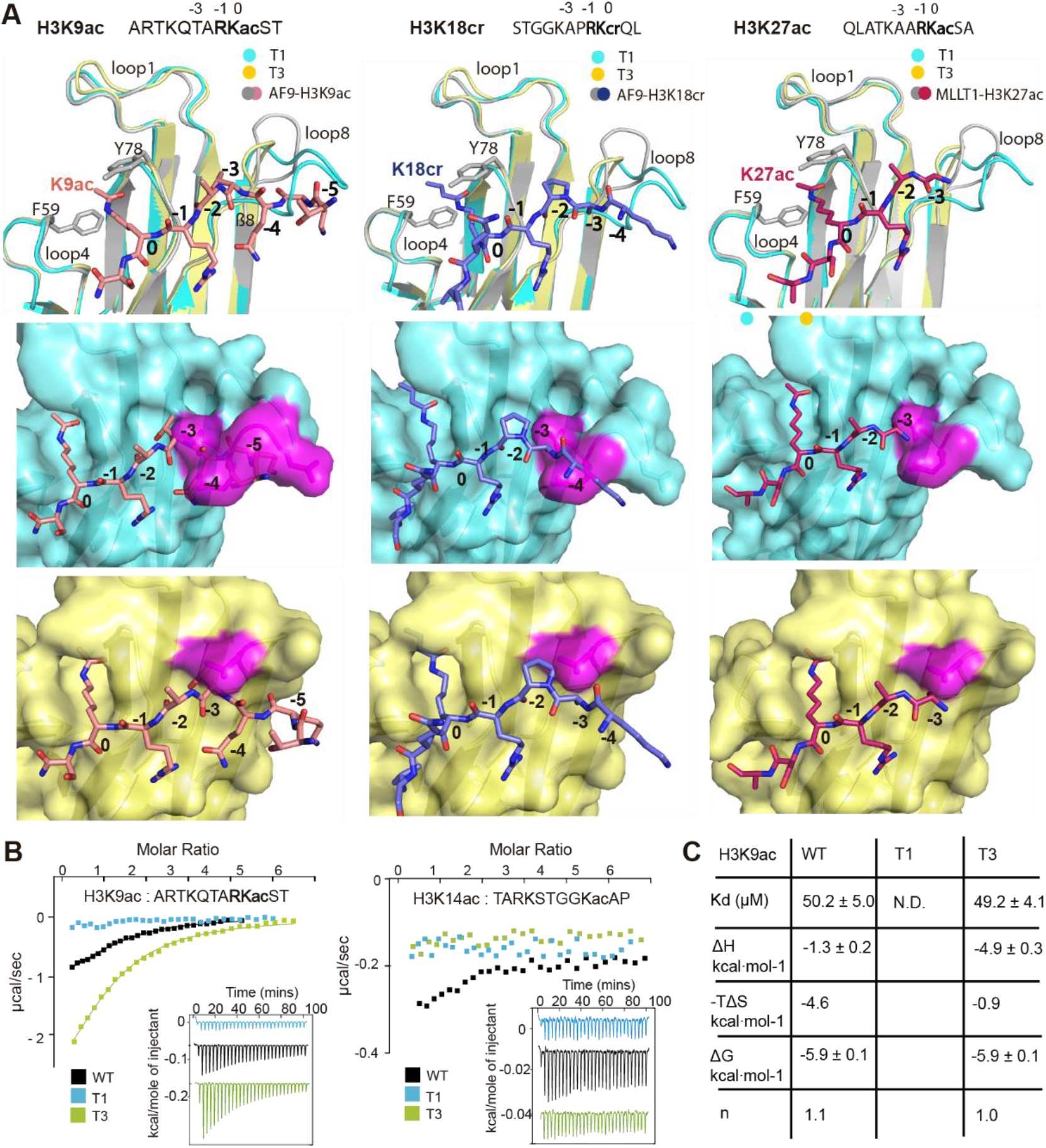
The oncogenic mutations cause an alteration of the binding interface for histone tail in MLLT1. (A) Superimposition between the AF9-H3K9ac (PDB ID: 4TMP), AF9-H3K18cr (PDB ID: 5HJD) and MLLT1-H3K27ac (PDB ID: 5J9S) complexes and T1 (cyan) and T3 (yellow) mutant structures revealed that the conformational changes of loop 8 in the mutants most likely perturbs the interaction with the N-terminal flanking region of these marks. Surface representations demonstrate an occlusion of the binding pocket in T1 mutant that are occupied by N-terminal residues of histone H3 in wild type MLLT1. In the T3 mutant this binding pocket is slightly impaired by the mutation. (B) Isothermal calorimetry (ITC) data measured on the binding of H3K9ac (left) and H3K14ac (right) in wild type and MLLT1 mutants. (C) The affinities and thermodynamic parameters from ITC for H3K9ac peptide; N.D. not determined. Because of the weak interaction, thermodynamic parameters have not been included for the T1 mutants.

Previous study by Wan et al. has demonstrated another unique characteristic of T1 mutant with an increased self-association that enhances its chromatin occupancy, leading to an increase in recruitment of super elongation complex and consequently aberrant transcription of several oncoproteins^*22*^. Although self-association in MLLT1 is likely driven mainly by the C-terminal intrinsic disordered region (IDR)^*22*^, we questioned whether the mutations in T1 might also play a role assisting oligomerization. Detailed analyses of the structures showed that MLLT1 YEATS domain formed a dimer in the crystals, which were generated by an application of crystallographic two-fold symmetry along the axis adjacent to β8 (Figure 3A, B and C). Although the dimeric assemblies of both mutants and wild type were mediated largely by the β8, the configuration and detailed interactions differed significantly. In wild type and T3, two protomers interacted in a head-to-tail fashion with their β8 strands arranged in an anti-parallel manner, enabling an inter-molecular β-sheet structure (Figure 3A, B and D). In contrast, T1 dimer was formed in a head-to-head manner with the β8 strands from both subunits associated in a parallel arrangement (Figure 3C). This unique orientation led to significant involvement of loop 8 in its dimerization. This loops from both protomers were paired in reverse, constructing large hydrophobic dimeric interface formed mainly by Phe105, Leu108, Pro112 and V114 (Figure 3E and F). Some inter-subunit hydrogen bond contacts were also observed, for example between loop 8 Asn111 and β8 His116, which for the latter is one of the insertion mutations, and loop 8 Asp103 and β8 His119, which for the latter had its position shifted along the β8 due to the mutation. In comparison, such contribution of loop 8 resulted in significantly larger dimeric interface in T1 mutant compared to wild type and T3, of which the shorter loop 8 in a different arrangement did not participate in oligomerization (surface buried area of 800, 293 and 329 Å^3^ for T1, T3 and wild type, respectively). However, we did not observe dimer formation of the mutated YEATS domains and wild type in solution (Figure 3G), which was rather expected since YEATS domain alone is unlikely to induce oligomeric assembly^*22, 24*^. Nevertheless, we postulate therefore that unique modulation of inter-molecular interface enabled by T1 insertion mutation may assist strong self-association characteristics of this mutant by promoting inter-molecular interactions between YEATS domains.

**Figure 3.**
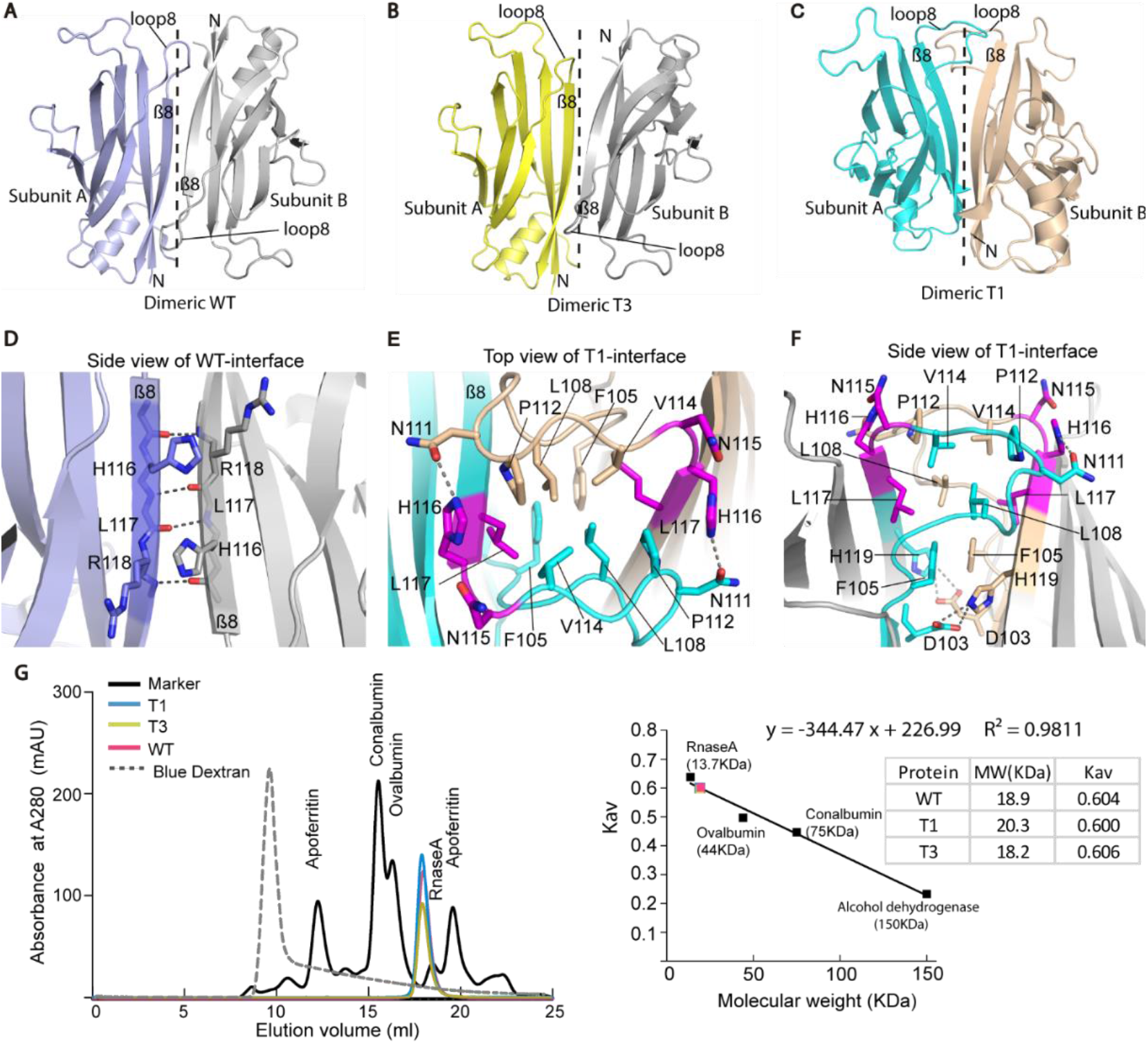
Comparison of dimeric assemblies of T1 and T3 mutants and MLLT1 wild-type in their crystal structures. Dimers of wild type (A, PDB ID: 6HQ0), T3 (B) and T1 (C) in the crystals generated by an application of crystallographic 2-fold symmetry along the β8 axis (dashed lines). (D) Close-up view of wild type dimeric interface at β8. (E, F) Dimeric interface of T1 with the insertion mutations colored in magenta. (G) Size exclusion chromatography elution profiles (left) and the plot of partition coefficients against molecular weight (right) demonstrate the monomeric behavior of the YEATS domains of both mutants and wild type (red dot) in solutions.

Previous reports have demonstrated that all oncogenic MLLT1 mutants still retains their acetyl/acyllysine reader function, which is required for their interaction with histone and their recruitment to chromatin^*22, 23*^. Thus, in a similar manner to wild type targeting the acetyl/acyllysine binding pocket of these mutants presents a strategy for inhibiting aberrant transcription of oncoproteins and hence the development of diseases. We have shown previously that the acetyl/acyllysine binding pocket of wild type is druggable^*17, 18*^, and recently a number of selective inhibitors have been developed, including a benzimidazole-amide-based derivative(**1**)^*18*^, SGC-iMLLT (**2**)^*19*^, NVS-MLLT-1 (**3**)^*21*^ and PFI-6 (**4**)^*20*^ (Figure 4A). To assess inhibitor binding in the mutants, we exploited isothermal calorimetry (ITC) to characterize the interactions between these inhibitors and both T1 and T3 mutants. As expected from the much unaltered acetyl/acyllysine binding pockets, the interactions for all inhibitors remained in both mutants with the affinities comparable to that observed for wild type (Figure 4B and C).

**Figure 4.**
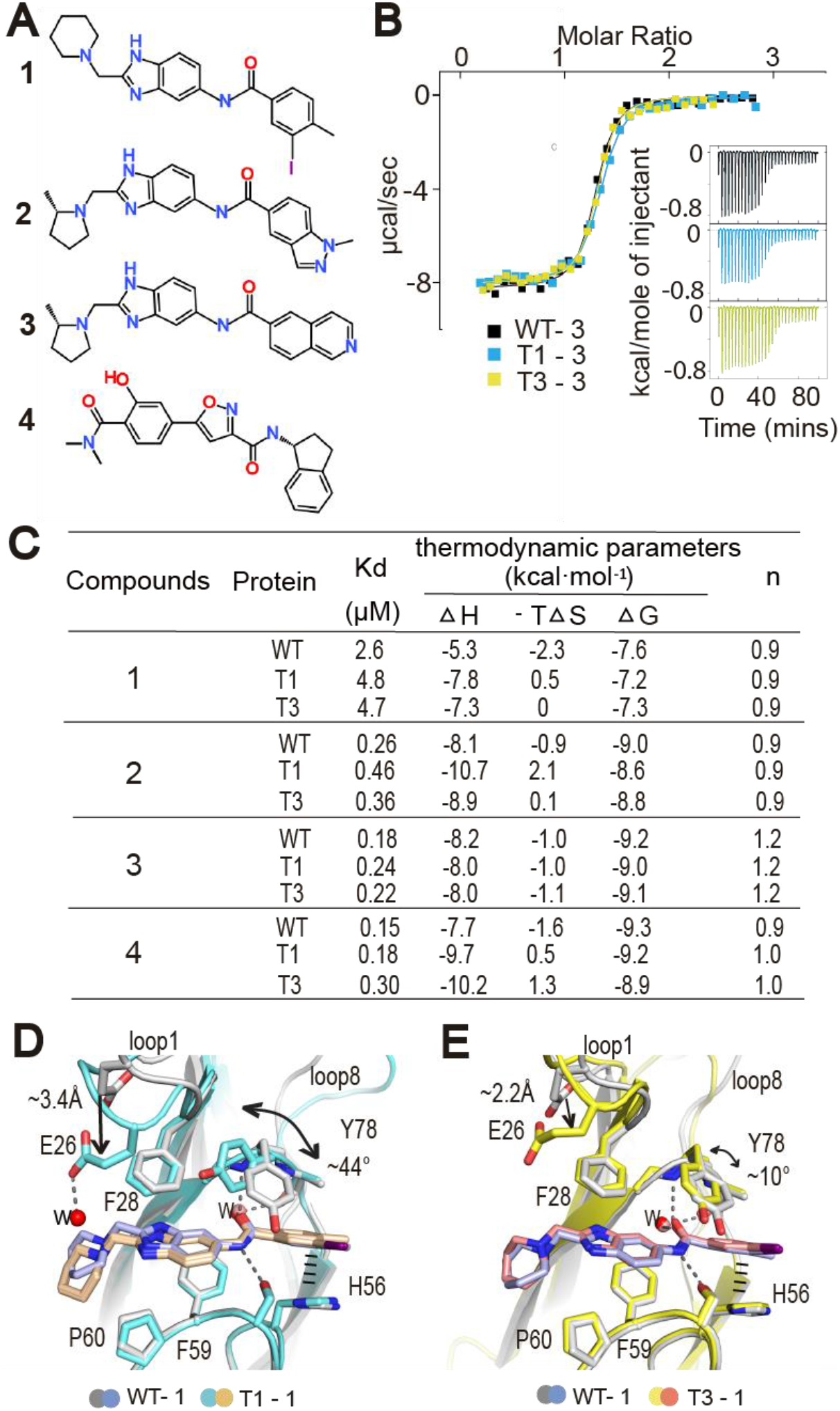
Analyses of inhibitor binding in T1 and T3 mutants. (A) Chemical structures of MLLT1 selective inhibitors, including benzimidazole-amide based compound 1, SGC-iMLLT (2), NVS-MLLT-1 (3) and PFI-6 (4). (B) Example of ITC integrated heat of binding for the interactions between T1 (cyan), T3 (yellow) and wild type (WT; black) with NVS-MLLT1-1 (3). Inset shows the isotherms of the raw titration heat. (C) Summary of ITC binding constants (KD) and thermodynamic parameters. (D and E) Structural comparisons of the binding of 1 between T1 and T3 mutants and wild type.

Gratifyingly, both T1 and T3 crystal structures were determined in complex with 1, offering therefore molecular insights into the inhibitor binding in these mutants. Structural comparison with the same inhibitor complex of wild type^*18*^ revealed that although the inhibitor adopted similar binding mode engaging fundamental key β-sheet-like hydrogen bond contacts to the backbone atoms of Ser76 and Tyr78, some differences between the mutants and wild type were evident (Figure 4D and E). Apart from various conformations of Tyr78 which has been shown to have high intrinsic flexibility^*18*^, loop 1 of the mutants, which is located in the proximity of loop 8 that contains the mutations, exerted a more closed conformation in contrast to an open form observed in wild type. This led to an alteration of Glu26 that moved towards inhibitor resulting in a more closed binding pocket. Such subtle structural changes were consistent with slight differences in thermodynamic signatures for the binding of most inhibitors in T1 and T3 with a trend of greater enthalpy opposed by slightly unfavorable entropy. However, further studies would be required to elucidate full mechanistic details how conformational change of loop 8 caused by the mutations affects loop 1 and in essence inhibitor binding properties. Nevertheless, our data suggested that both MLLT1 mutants and wild type could be targeted by similar inhibitors, yet small alterations of inhibitor binding properties might be expected potentially due to an indirect influence of the mutations located within the second sphere of the binding site.

## Conclusion

The mutations in YEATS domain have recently been discovered as a factor that caused aberrant function of MLLT1 driving development of aggressive cancer. Our data demonstrated the structural consequences of both insertion and substitution with deletion mutations induces significant conformational changes in loop 8. Thus, the data presented here provide a structural basis for their altered activities for acetylated/acylated histone binding due to perturbation of the binding interface. In addition, unique dimeric assembly of T1 observed in the crystals was likely induced by the mutated loop 8, offering a potential explanation on the proposed contribution of the mutation in YEATS domain towards an increased self-association of this MLLT1 mutant that leads to continuous activation of transcription of oncogenes^*22*^. Inhibitor binding studies suggested that the mutants can generally be targeted by the developed chemical probes that have been designed for the wild type protein. However, due to the structural alterations cause by the mutations, it might be possible designing mutant specific inhibitors. Together, our structural insights provided insights into the molecular consequences caused by MLLT1 mutations perturbing the reader function of this important epigenetics reader. In addition, since the acetyl/acyllysine reader activity remains crucial for maintaining cell homeostasis, targeting these MLLT1 mutant forms with small molecules presents an attractive strategy for the development of new treatment strategies in MLLT1 mutated cancer.

## EXPERIMENTAL SECTION

### Protein expression and purification

Wild type MLLT1 YEATS domain (aa. 1-148) was subcloned into pNIC-CH. Both T1 and T3 mutants were generated by PCR-based site-directed mutagenesis. All proteins harbouring C-terminal-His_6_-tag were overexpressed in *E. coli* Rosetta. Cells were initially grown at 37 °C until OD_600_ reaching 1.6-1.8, and subsequently cooled to 18 °C and at the OD_600_ of ∼2.6-2.8 induced with 0.5 mM IPTG overnight. The recombinant proteins were purified initially by Ni^2+^-affinity chromatography and subsequently by size exclusion chromatography in buffer containing 25 mM Tris pH 7.5, 300 mM NaCl, 0.2 mM TCEP, 5% (v/v) glycerol.

### Crystallization, Data collection and Structure determination

Recombinant T1 and T3 proteins were concentrated to ∼9 mg ml^-1^ and incubated with compound 1 at 4 °C for one hour before crystallization. The crystals of both mutants were obtained by sitting drop vapor diffusion method at 20 °C. T1 crystals grew in the condition containing 25% (w/v) PEG3350, 0.1 M bis-tris, pH 5.5, while T3 crystals were obtained from 28% (w/v) PEG3350, 0.2 M sodium acetate trihydrate, 0.1 M bis-tris, pH6.0. All crystals were cryoprotected with mother liquor supplemented with 20-22% (v/v) ethylene glycol before flash cooled in liquid nitrogen. Diffraction data were collected at Swiss Light Source, X06SA. Data were processed with XDS^*25*^ and scaled with aimless^*26*^. Structures were initially solved by molecular replacement using PHASER^*27*^ and the structure of wild type MLLT1 (PDB ID: 6T1L). COOT^*28*^ was used for model rebuilding and REFMAC^*29*^ for refinement. The final models were verified for geometry correctness with Molprobity^*30*^. Data collection and refinement statistics are summarized in Table 1.

### Isothermal titration calorimetry

Compounds used in this study were from previous studies^*18, 19*^ or obtained from the chemical probe program of the Structural Genomics Consortium ^*20, 21*^. Isothermal titration calorimetry (ITC) experiments were performed using NanoITC instrument (TA Instrument) at 25 °C in buffer containing 25 mM Tris, pH 7.5, 300 mM NaCl, 0.2 mM TCEP and 5% (v/v) glycerol. Proteins at 200-300 µM were titrated into 20-30 µM inhibitors. Data analysis was performed with NanoAnalyze™ Software (TA Instruments) using an independent binding model.

## Accession codes

Coordinates and structure factors have been deposited with accession codes 7b10 and 7b0t.

## AUTHOR INFORMATION

## ACKNOWLEDGMENT

The SGC is a registered charity that receives funds from; AbbVie, Bayer AG, Boehringer Ingelheim, Canada Foundation for Innovation, Eshelman Institute for Innovation, Genentech, Genome Canada through Ontario Genomics Institute [OGI-196], EU/EFPIA/OICR/McGill/KTH/Diamond, Innovative Medicines Initiative 2 Joint Undertaking [EUbOPEN grant 875510], Janssen, Merck KGaA (aka EMD in Canada and US), Merck & Co (aka MSD outside Canada and US), Pfizer, São Paulo Research Foundation-FAPESP, Takeda and Wellcome. The authors thank staff at SLS for their support during crystallographic X-ray diffraction testing and data collection and the Frankfurt Cancer Institute as well as the German translational cancer network (DKTK) for support.

## ABBREVIATIONS

YEATS domain: The Yaf9, ENL, AF9, Taf14, Sas5 (YEATS) domain;
ENL: eleven-nineteen-leukemia protein;
MLLT1: myeloid/lymphoid or mixed-lineage leukemia translocated to, chromosome 1 protein;
AF9: ALL1-fused gene from chromosome 9 protein;
MLLT3: myeloid/lymphoid or mixed-lineage leukemia translocated to chromosome 3 protein.

